# Comprehensive healthspan assessments and influence of sex as a biological variable in aging rats

**DOI:** 10.1101/2025.09.22.677661

**Authors:** Abhishek Chandra, Zaira Aversa, Xu Zhang, Joshua N. Farr, Achudhan David, Ritika Sharma, Tamara Tchkonia, James L. Kirkland, Paul D. Robbins, Laura J. Niedernhofer, Diana Jurk, João F. Passos, David G. Monroe, Jordan Miller, Nathan K. LeBrasseur, Yuji Ikeno, Sundeep Khosla, Robert J. Pignolo

## Abstract

Rats share a significant amount of genetic and physiological similarity with humans. Many biological processes and pathways are conserved between rats and humans, making rats a suitable model for studying various aspects of human health and disease. Using rats as an aging model offers a more ethical alternative to using larger, longer-lived animals like primates. Rats are easier to handle in laboratory settings, as compared to non-human primates, both of which have physiological functions like humans. To date, there are very few studies which have comprehensively studied age-related changes in rat physiology. Here we present a longitudinal assessment of several aspects of Brown-Norway rat physiology and histopathology using molecular and functional assessments at 6-, 17- and 27 months of age. Our studies thus provide age-related healthspan parameters, which can be used as reference for genetic or pharmacological rat models of aging.

## Introduction

With improved diet, lifestyle, and advances in medical sciences, there is an increase in the aging population throughout the world. With increasing age, age-related morbidities are also on the rise [1], and while modern medicine strikes to tackle many of these morbidities efficiently, the hope remains to develop improved therapeutics for single or multiple co-morbidities, ideally by addressing fundamental aging processes that underly multimorbidity.

Rats, like humans, experience various age-related changes in their tissues, organs, and overall physiology. These changes can include reduced physical activity, cognitive decline, increased susceptibility to diseases, and alterations in metabolism. Rats are used to model a wide range of diseases, including cardiovascular diseases [2], diabetes, cancer, neurological disorders [3], and more, and for certain disease states are superior to mice in recapitulating the human conditions [4]. By inducing specific disease conditions or genetically modifying rats to mimic human diseases, researchers can study disease mechanisms, test potential therapies, and explore disease progression.

Mice are among a few preferred models for studying physiological functions and disease states. This is largely because of the availability of a greater number of genetic models in mice than in rats. In more recent times more rat transgenic models have become available to better understand pathological diseases [5–8]. This expansion of rat transgenic models offers researchers new avenues to investigate complex diseases and physiological processes with greater specificity and relevance to humans. Many FDA-approved drugs have gone through testing in rat models due to a lack of more appropriate models. The use of non-human primates has declined due to ethical concerns, and rats have remained the model of choice for preclinical testing, as a prelude to clinical trials in humans [9, 10].

Very few studies have performed comprehensive health assessments of the Brown Norway rat. Here we provide a rigorous longitudinal assessment of molecular, physiological, and functional parameters at different ages. Age-related phenotypes and characteristic physiologic aging hallmarks presented in this comprehensive analysis provides the basis for comparisons to genetic or pharmacological manipulations done using the Brown-Norway rat model.

## Results

### Frailty and physical functioning

The frailty index (FI)[11] [12] score was performed as described previously [13]. Mean FI scores are presented which is based on a 30-parameter scoring system [13]. The FI score progressed with age in both males and females (Fig. 1A). However, these changes were much more pronounced in males where FI scores were significantly higher in 17-month-old males, while there was no change in the FI score of 17-month-old females. FI score in 27-month-old rats showed no difference in both males and females when compared to 6-month-old rats (Fig.1A).

**Figure 1.**
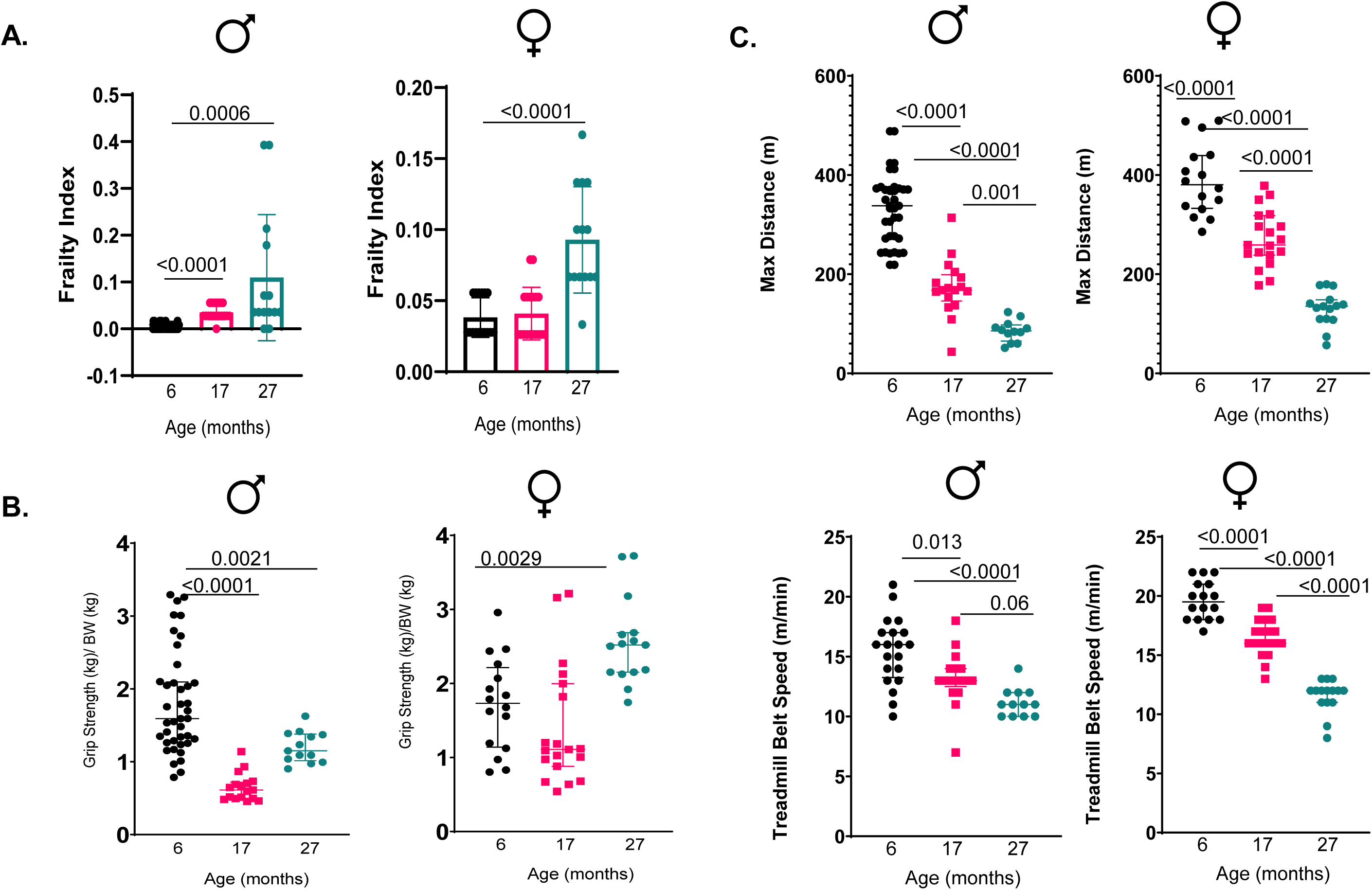
Age-related frailty scoring and physical performance. (A) Individual FI scores for 6-, 17- and 27-month-old male and female rats are displayed as dot plots. The frailty measures which were based on 30 parameters, ranged from assessing the integument, physical or musculoskeletal signs, auditory changes, ocular and nasal observations, digestive and urogenital changes, respiratory rates and discomfort. (B) Grip strength measurements are presented in 6-, 17- and 27-month-old male and female rats with normalizations made to the body weight. (C) Physical performance was measured using a treadmill test in which maximum distance traveled, and highest treadmill belt speed reached, was measured in the 6-, 17- and 27- month-old male and female rats. All statistical measurements are done using One-way ANOVA.

To assess muscle function, a grip strength test was used. As expected, aged male rats performed poorly on this test and had lower scores as compared to young rats (Fig. 1B). Conversely and surprisingly, female rats at 17-month did not show any difference in the grip strength parameter, while 27-month-old females, showed an increased grip strength (Fig. 1B). Next, we assessed the ability of these rats to run on a treadmill. Using maximum distance ran and an arbitrary treadmill belt speed, both male and female 27-month-old rats were the worst performers as compared to both 6-month and 17-month-old rats, while 17-month-old rats performed worse than their 6-month-old counterparts (Fig.1C).

### Voluntary movement and threat avoidance

Thigmotaxis refers to the tendency of animals to stay close to the walls or periphery of an open area[14]. In the open field test, anxious rats tend to exhibit increased thigmotaxis, spending more time along the walls rather than exploring the center of the open field. This behavior suggests an attempt to seek safety in the less exposed areas. We observed that aged rats tend to spend less time in the central, more exposed area of the open field compared to the periphery (Fig. 2). This preference for the periphery is thought to be related to their desire to avoid potential threats. We measured the total distance traveled as well as the total distance traveled to the center and observed that both parameters declined with age.

**Figure 2.**
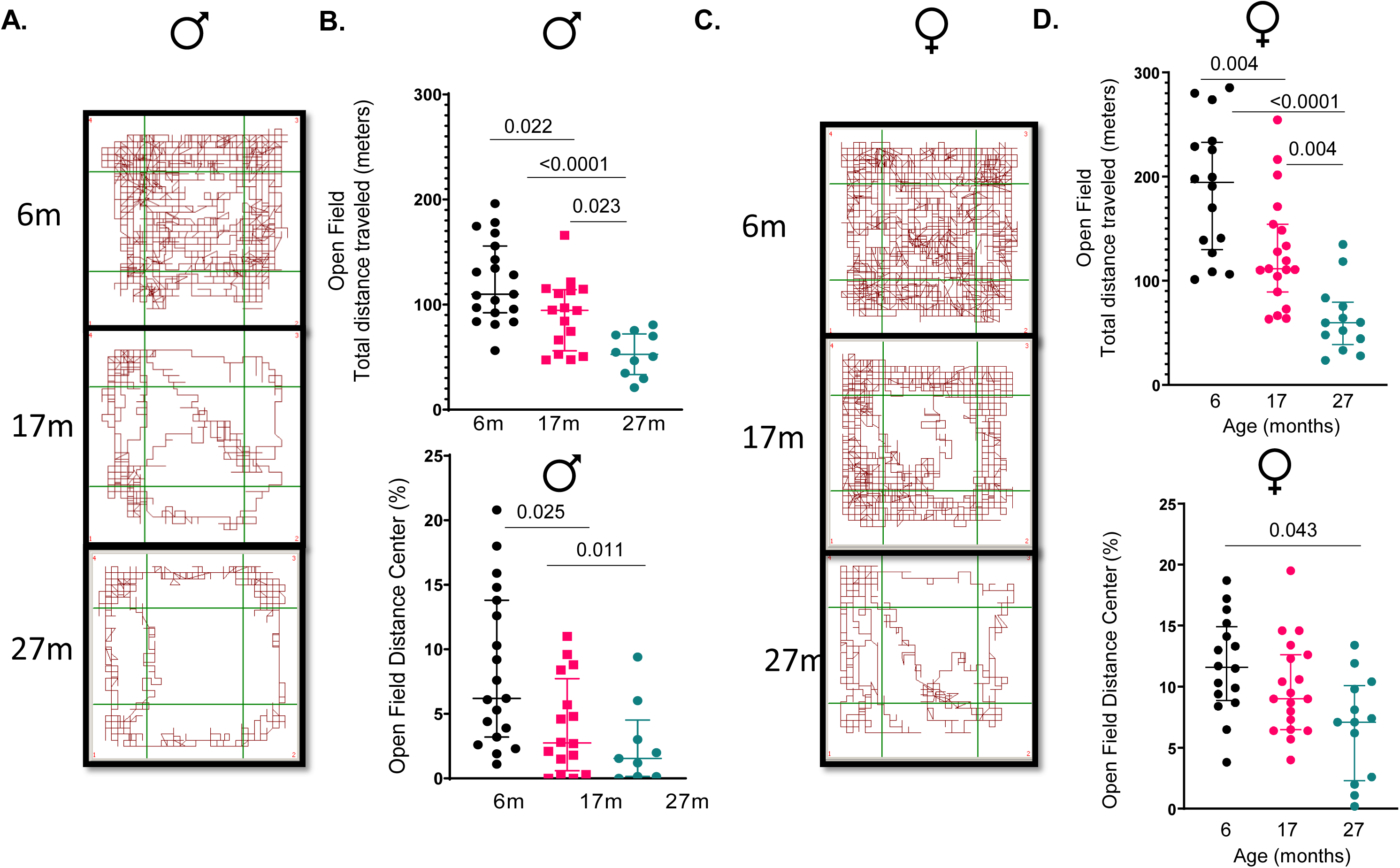
Age-related changes in anxiety measurements using Open Field Test. The images represent the movement of the 6-, 17- and 27-month-old male (A) and female (C) rats in the open field chamber. Total distance traveled (B, D, top panel) or distance traveled to the center (B, D, bottom panel), during the time spent in the chamber is shown in 6-, 17- and 27- month-old male (B) and female (D) rats. All statistical measurements are done using One-way ANOVA.

### Changes in body fat and glucose tolerance

Body fat in obesity has been correlated negatively with glucose tolerance. Our data shows that body fat increases in male rats in age-dependent manner, while the female rats tend to have higher weights at 17 months, but not at 27 months. The 27-month-old rats which had the highest weights (Fig. 3A), also showed the least glucose tolerance (Fig. 3B) when compared with younger rats. As compared to male rats, glucose tolerance in females was unaffected by increasing age.

**Figure 3.**
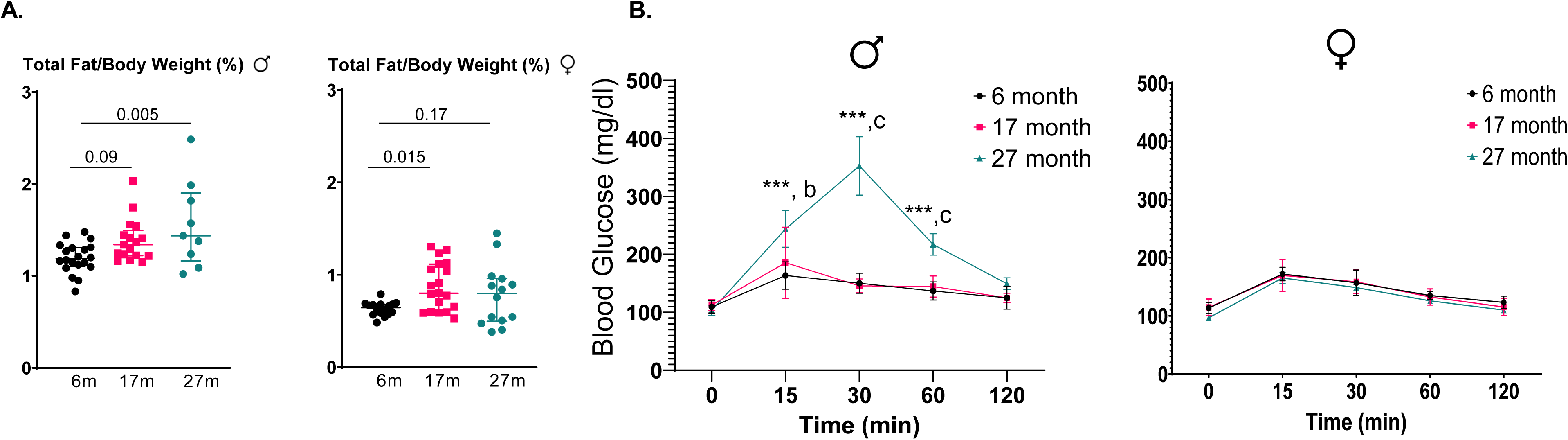
Age-related changes in body fat and glucose tolerance. The graphs represent the wet weight for total fat normalized to the body weights in the 6-, 17- and 27-month-old male and female rats (A). (B) Glucose tolerance test (GTT) measurements using 2μl of blood with glucose being measured using a handheld glucometer at time 0, 15-, 30-, 60- and 120-min after the rats were fasted and fed with glucose. All statistical measurements are done using One-way ANOVA.

### Anatomical and molecular changes in rat muscle and bone

One of the most noticeable changes in aging rats is the loss of muscle mass, a condition known as muscle atrophy or sarcopenia. This occurs due to a decrease in the size and number of muscle fibers. Age-related muscle atrophy is often attributed to a reduction in muscle protein synthesis, an increase in muscle protein degradation, and changes in hormonal profiles. As shown in Fig.1C muscle function declined with age when assessed by treadmill testing. Similarly, declines in muscle mass were observed in the quadriceps femoris, gastrocnemius, and soleus. These declines in muscle mass, when normalized to body weight, was significantly lower in 17-month- and 27- month-old female and 17-month-old male rats (Supp. Fig. 1). Due to strict COVID-19 restrictions in 2020 and reduced availability of staff, the 27-month-old males were not assessed for muscle mass.

Markers of cellular senescence are molecular or cellular features that indicate the onset or presence of a state of irreversible growth arrest accompanied by the senescence-associated secretory phenotype (SASP). Cellular senescence is a natural biological process that contributes to various aspects of aging, tissue repair, and the prevention of proliferation of damaged or potentially cancerous cells. Cyclin-dependent kinase inhibitors (CDKIs), CDKN1A or p21, and CDKN2A or p16^INK4a^ are markers of cellular senescence. While femurs from 27-month-old male and female mice showed a 2-3-fold increase in *p16^Ink4a^* expression, there was no change in the *p21* transcript (Fig. 4A & 4B). Conversely, muscle showed elevated levels of both *p16^Ink4a^*(2-3-fold change in 27-month-old male and female mice vs 6-month-old mice) and *p21* (3-7-fold change in 27-month-old male and female mice vs 6-month-old mice) (Fig. 4A & 4B). Similar increases in *p21* and *p16^Ink4a^*transcripts were seen in inguinal fat in 27-month-old males (Fig. 4A). Interestingly, there were no changes in either senescence marker in inguinal fat from female rats (Fig. 4B).

**Figure 4.**
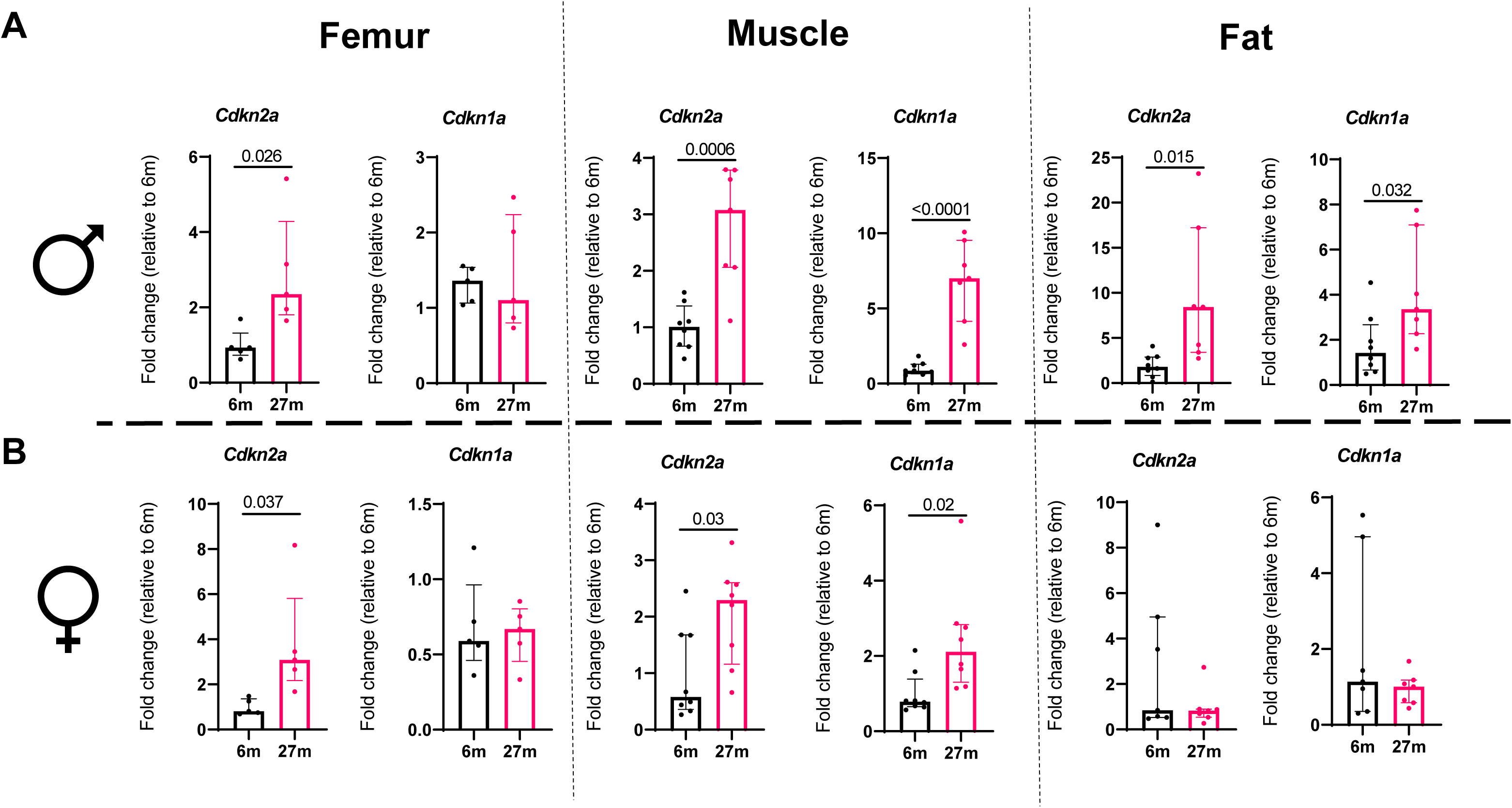
Age-related changes in gene expression for markers of senescence. Gene expression changes for *Cdkn2a* (*p16^Ink4a^*) and *Cdkn1a* (*p21*) were detected in femur, muscle and fat tissues from 6- and 27-month-old male and female Brown Norway rats. Statistical analyses were done using GraphPad Prism and p values were calculated using two-tailed unpaired t-tests.

CD68 is highly expressed in monocytes, circulating macrophages, and tissue macrophages. We observed an increase in the *Cd68* transcript in 27-month-old male and female rats as compared to younger rats (Fig. 5A, B). Among the SASP factors we analyzed, *Cxcl10* was significantly elevated in skeletal muscle in older rats in both males and females, while *Tnf* showed a trend toward increasing in aged male rats but showed a significant increase in expression in older female rats both at 17- and 27 months.

**Figure 5.**
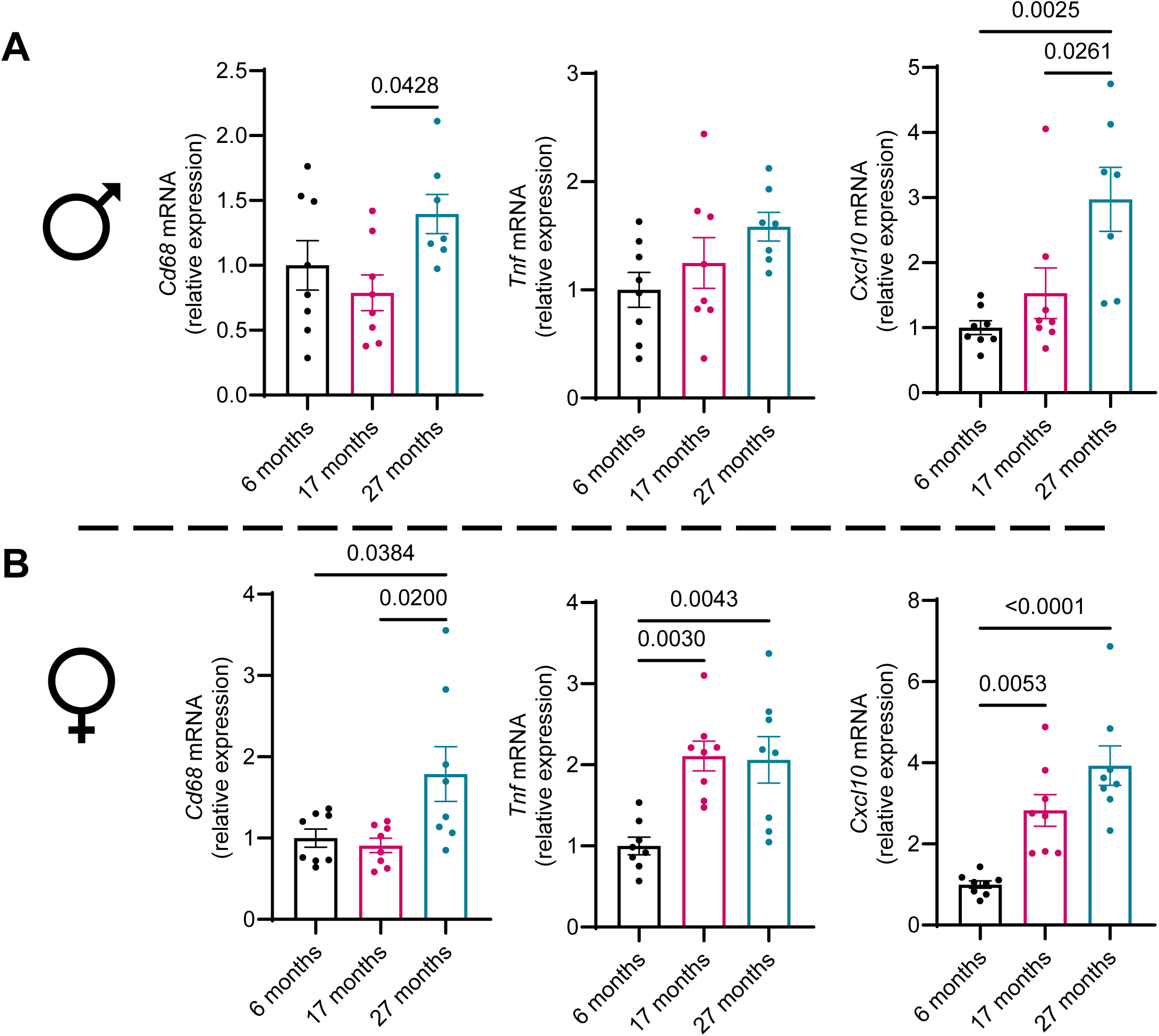
Age-related changes in gene expression for muscle specific SASP genes. Gene expression changes for SASP markers such as *Tnf* and *Cxcl10* were measured in muscle tissue from 6-, 17- and 27-month-old male and female Brown Norway rats. Statistical analyses were done using GraphPad Prism and p values were calculated using One-way ANOVA.

Analyzing bone parameters, we observed a decline in spine bone volume fraction in both 17- and 27-month-old male rats (Fig.6A). The male rats reached their nadir of spine BV/TV by 17 months with no further decline seen in 27-month-old rats. Interestingly, spine BV/TV declined in 17-month-old female rats, but showed reversal by 27-months. (Fig. 6G). Spine trabecular numbers (Tb.N) declined in both male and female aged rats at both 17- and 27-months, while an age-related increase was observed in both trabecular thickness (Tb.Th) and trabecular separation (Tb.Sp) (Supp. Fig.2A). Consistently, similar trabecular bone loss was observed in the femora in both male and female rats in an age-dependent manner (Fig. 6B, 6H). Similar to the spine, the femoral Tb.N decreased significantly at both 17- and 27 months in both male- and female-rats. Interestingly, and contradictory to spine data, the Tb.Th declined in both male- and female-rats at 27 months, while the Tb.Sp showed a similar increase with age as was seen in the spine (Supp. Fig. 2B). No significant changes were observed in femoral cortical thickness. While femoral cortical volumetric bone mineral density (vBMD) increased only in 27-month-old male rats (Fig. 6D), there was an age-dependent increase in vBMD of femoral cortical bone in female rats (Fig. 6J). Conversely, cortical porosity declined in male rats in an age dependent manner (Fig. 6E), with no significant changes seen in aged female rats. We next evaluated the failure to load using finite element analysis (FEA) and observed a significant decline in both 27-month-old males (Fig.6F) and in 17- and 27-month-old females (Fig.6L).

**Figure 6.**
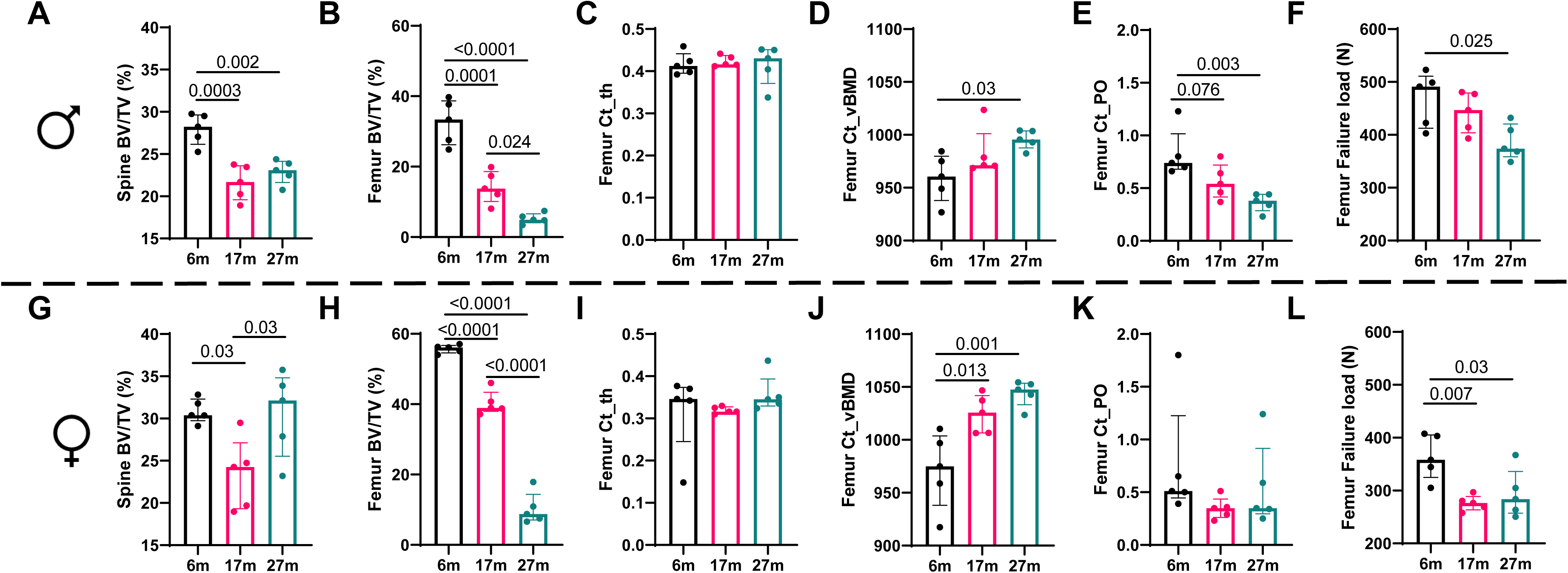
Age-related changes in bone architecture in rats. Micro-CT scans were done to analyze bone volume fraction in spine (A, G) and femur (B, H), cortical thickness (Ct_th) (C, I), cortical volumetric bone mineral density (Ct_vBMD) (D, J) and cortical porosity (Ct_PO) (E, K), and failure to load as detected by finite element analysis (F, L) are shown in 6-, 17- and 27-month-old male and female Brown Norway rats. Statistical analyses were done using GraphPad Prism and p values were calculated using One-way ANOVA.

### Changes in complete blood cell counts

Significant declines were observed in the total white blood cells (WBCs) and lymphocytes in both aged male and aged female rats, when compared to the young mice, however the numbers largely remained within the threshold limit, with aged female rats, having lower WBC and lymphocyte counts than the normal range (Fig.7). While the monocyte counts remained within the normal range, there was a significant decline in the monocyte numbers in 27-month-old female rats, when compared to young female rats, with no change seen in male rats. No change was observed in the neutrophil counts, however, there was a strong trend of decline in female rats. When sex comparisons were done at 27 months, significantly lower counts of WBCs, lymphocytes, and neutrophils were observed in aged female rats, with no change seen in monocyte counts (Fig. 7C). The 27-month-old female rats were more anemic as compared to young rats, with significant declines in RBC numbers, with decline in hemoglobin levels (Fig. 8). While there was a decline in RBC counts in aged male rats, both RBC and hemoglobin stayed within the normal range. Hematocrit is a blood test that measures the proportion of red blood cells in the blood, and lower hematocrit count in 27-month-old female rats indicate anemia as compared to their 6-month-old counterparts, however, lower hematocrit values in aged male rats were largely within the normal range (Fig. 8). Platelet, which is one of the components of a clot, was reduced in both male and female rats (27-months vs 6-months) and lower than normal range as well (Fig.8). Aged females were generally more anemic than aged males and had reduced ability to form clots as well (Fig. 8).

**Figure 7.**
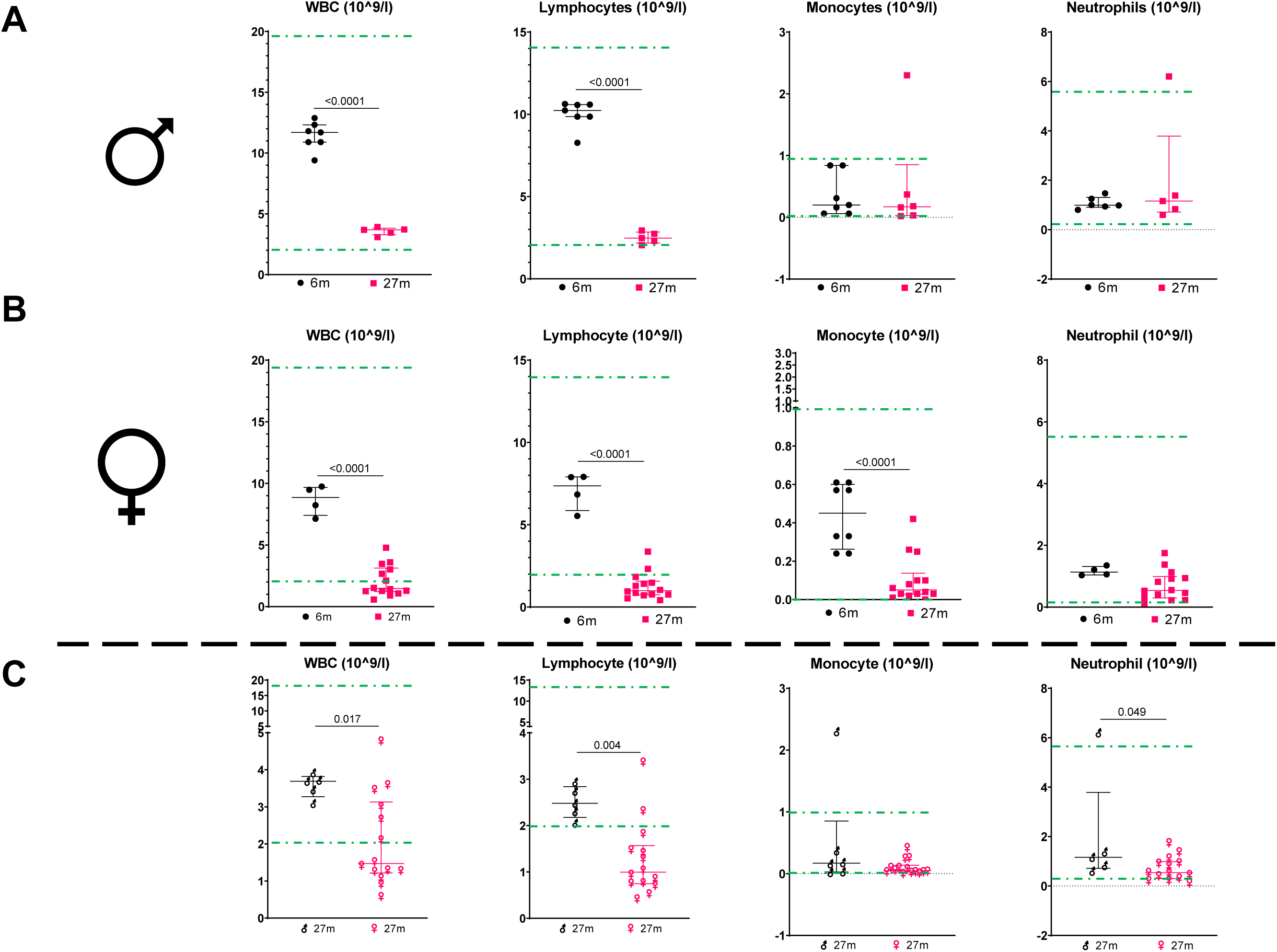
Age-related changes in white blood cells. (A, B) Whole blood counts were performed on the blood from 6- and 27-month-old male and female Brown Norway rats, and number of white blood cells (WBCs), lymphocytes, monocytes and neutrophils are reported here. (C) Comparison of cell counts between male and female 27- month-old rats is shown using symbols for male (♂) and female (♀). Statistical analyses were done using GraphPad Prism and p values were calculated using two-tailed unpaired t-tests.

**Figure 8.**
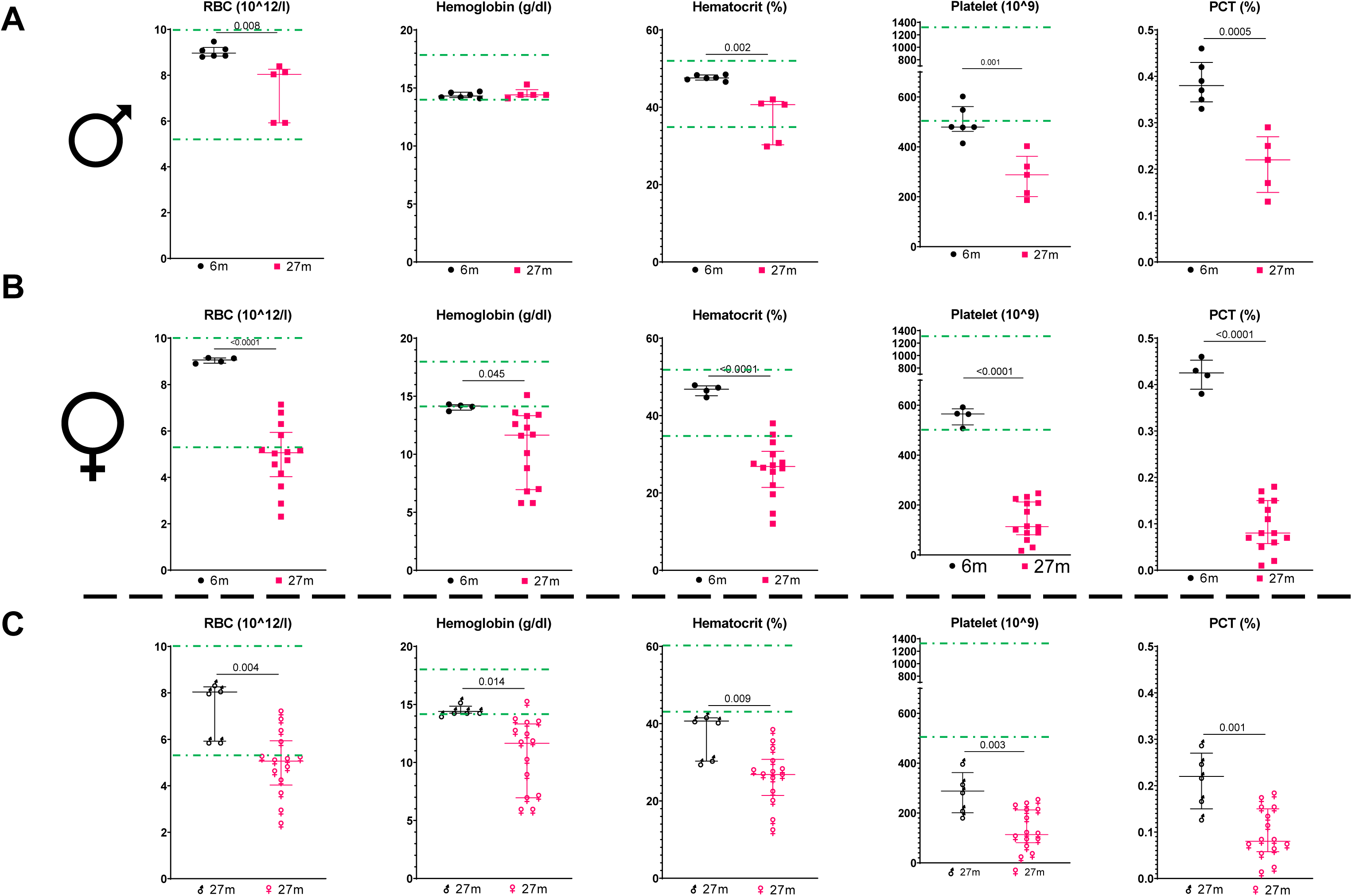
Age-related changes in red blood cells and hemoglobin. (A, B) Whole blood counts were performed on the blood from 6- and 27-month-old male and female Brown Norway rats, and number of red blood cells (RBCs), hemoglobin count, percentage hematocrit level, platelet count and PCT as the volume occupied by platelets in the blood as a percentage are reported here. (C) Comparison of RBC, hemoglobin, hematocrit percentage, platelet count and PCT percentage between male and female 27-month-old rats is shown using symbols for male (♂) and female (♀). Statistical analyses were done using GraphPad Prism and p values were calculated using two-tailed unpaired t-tests.

### Changes in liver and kidney function

Alanine aminotransferase [17] is an enzyme found primarily in the liver. It plays a role in converting proteins into energy for the liver cells. Aspartate aminotransferase (AST) is an enzyme that plays a key role in amino acid metabolism. It is found in various tissues throughout the body, with high concentrations in the liver, heart, muscles, kidneys, and brain. Both are indicators of liver function and disease. Both ALT and AST were significantly elevated above the normal range in 27-month-old male rats compared to younger rats, while the increased levels in aged female rats reached significance for AST but not for ALT (Fig.9). Interestingly, 27-month-old female rats had lower ALT and AST levels as compared to the males.

**Figure 9.**
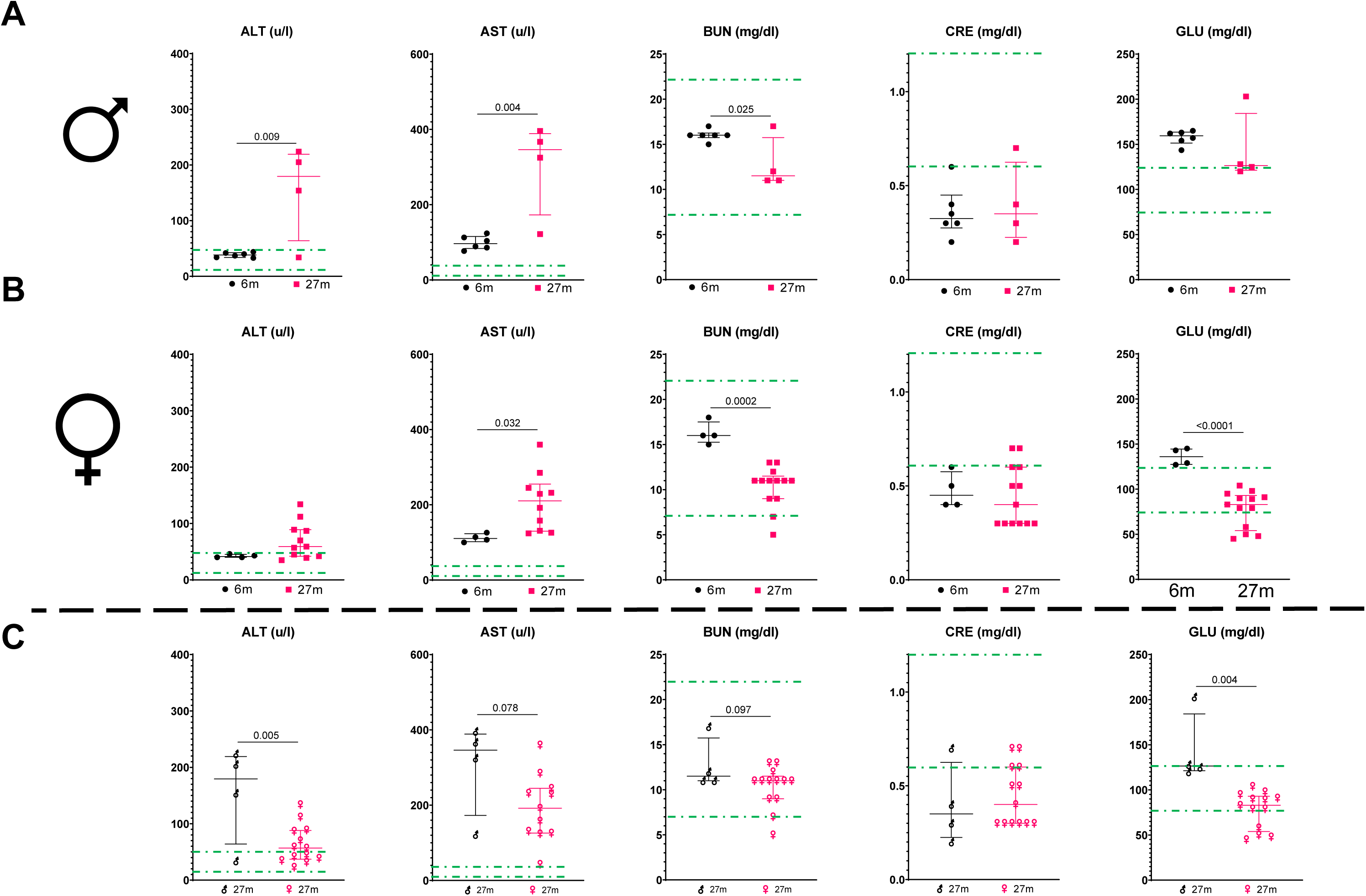
Age-related changes in liver and kidney function markers. (A, B) Liver and kidney function tests were performed on the blood from 6- and 27-month-old male and female Brown Norway rats. Alanine aminotransferase (ALT) and Aspartate aminotransferase (AST) were used for liver function, Blood urea nitrogen (BUN) levels and serum [52] creatinine (CRE) test (a blood test used to check how well your kidneys are filtering your blood), and random blood glucose was measured in 6- and 27-month-old male and female rats. (C) Comparison of liver and kidney function tests was made between male and female 27-month-old rats, is shown using symbols for male (♂) and female (♀). Statistical analyses were done using GraphPad Prism and p values were calculated using two-tailed unpaired t-tests.

Blood Urea Nitrogen (BUN) and creatinine (CRE) are commonly measured parameters in blood tests related to kidney function. Both male and female rats at 27 months had lower BUN values compared to younger animals, with females having a lower BUN than age-matched males at 27 months, however, the BUN values were mostly within the normal range (Fig.9). Surprisingly, the creatinine levels were lower than normal range at all ages with no change with sexes. No change in creatinine levels was observed with old age as compared to young rats.

### Cardiac function with age

The diameter of the heart chambers, such as the left ventricle, plays a critical role in cardiac function. The diameter affects the volume of blood the chamber can hold and in part subsequently limits the percent volume that can be ejected during systole (contraction) and filled during diastole (relaxation). Changes in chamber diameter can impact stroke volume, which is the amount of blood pumped with each heartbeat. Passive pressure, also known as preload, is the pressure exerted on the walls of the heart chambers when they are filled with blood. This pressure is a critical determinant of the heart’s ability to stretch and fill during diastole. It is related to the volume of blood returning to the heart (venous return) and the compliance (elasticity) of the heart muscle. When passive pressure was plotted against diameter, there were no significant differences observed amongst all three age groups (Supp. Fig. 3A). Aging can lead to decreased compliance of the heart muscle, making the heart less effective at pumping blood. Indeed, the 27-month-old rats showed reduced compliance at lower pressure values (Supp. Fig. 3B). It is expected that with old age, the distensibility of the heart may decrease, which can affect the heart’s ability to adapt to changing blood volume. As compared to 6-month-old rats the 17-month-old and 27-month-old rats showed a trend toward a decline in their distensibility (Supp. Fig. 3C).

Lower heart weights were noted at all ages in females as compared to males, but this was proportional to total body weight differences between the sexes (Supp. Fig. 3D-F). When both sexes were grouped together, the heart weight increased with age (Supp. Fig. 3E, F).

### Histopathological evaluation

Nephropathy indicates a deterioration of kidney function. Both male and female rats show a significant increase in nephropathy grade with age (Fig.10). For males, the grade is significantly higher at 17- and 27-months, compared to 6-month-old rats (Fig.10A). In females, a similar pattern is observed, with significant increases at both 17- and 27-months, compared to 6-month-old female rats (Fig. 10B). An inflammation score is a pathology score that summarizes the degree of inflammation in a tissue sample. In both male and female rats, the inflammation score increases significantly from 6 months to 27 months but does not increase until 17 months (Fig.10). Increase in fat content in cells is a sign of tissue dysfunction. A marked increase in fatty change is seen at 27 months in male rats. In females, fatty change increases significantly at 17 months, while showing a trend to increase at 27 months (Fig.10). Disease burden was also scored in both male and female rats. While females experience an increase in overall disease burden at 17- and 27- months of age, the male rats showed no change throughout the age (Fig.10).

**Figure 10.**
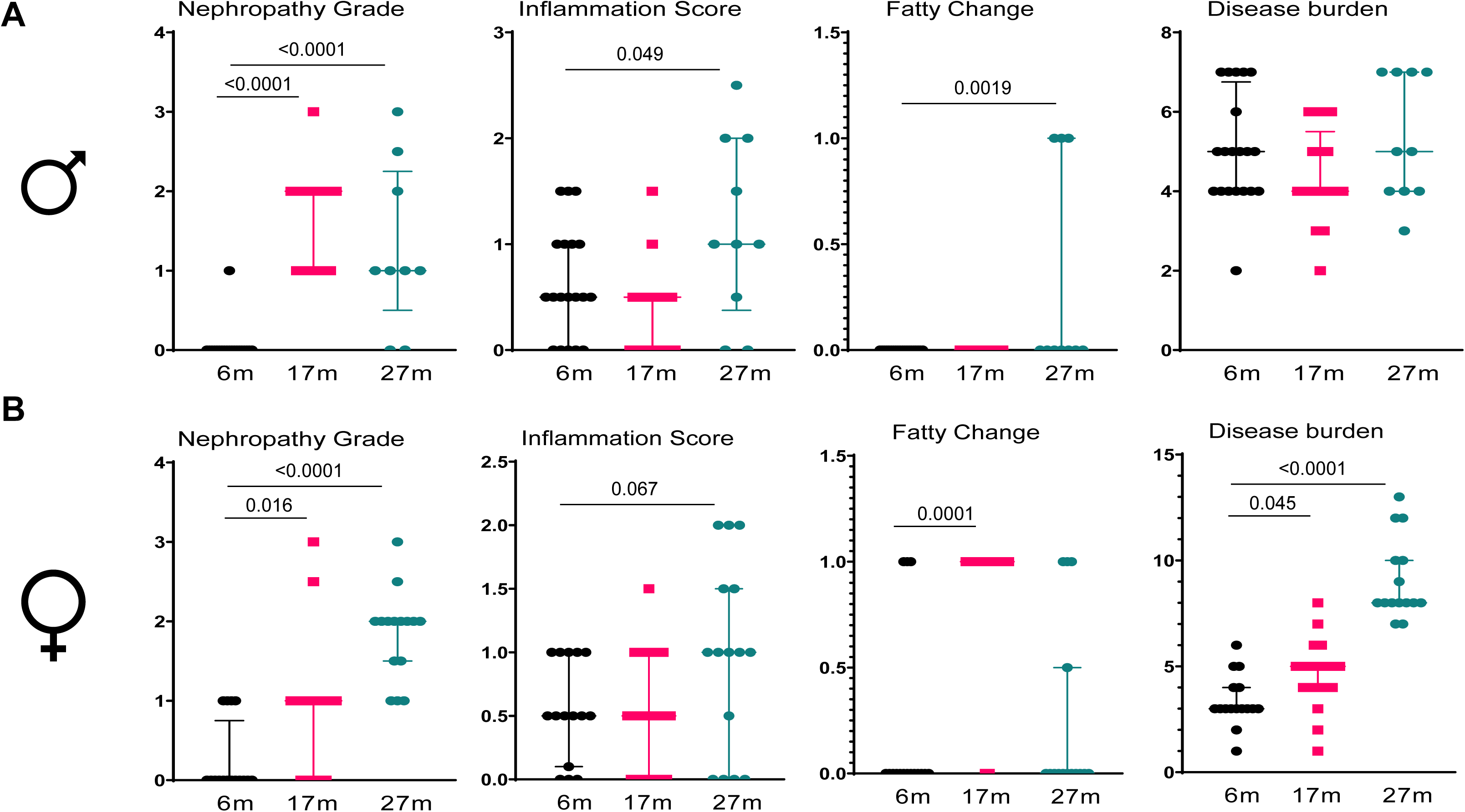
Age-related histopathological changes. The figure represents various histopathological parameters including nephropathy, inflammation score, fatty change and overall disease burden. Comparisons were made between male (♂) (A) and female (♀) (B) rats of different ages. Statistical analyses were done using GraphPad Prism and p values were calculated using One-way ANOVA.

## Discussion

The world’s population is experiencing a significant increase in the proportion of older adults. This demographic shift is driven by factors such as declining fertility rates, improvements in healthcare and living standards, and longer life expectancy[18]. The increasing aging population presents unique challenges, with increasing multi-morbidities and often reduced quality of life for older individuals. It has thus become important to understand the physiology of aging tissues and organ systems. Preclinical studies in rodents are considered a prerequisite before a clinical trial and a potential approval by regulatory agencies for a particular therapeutic or surgical procedure. Several parameters associated with aging are not well understood. Our study identifies molecular, physiological, and functional parameters that track with normal aging and together describe much of the aging phenotype in Brown-Norway rats. A subset of these parameters illustrates the importance of sex as a biological variable in the aging process.

One of the most common observations in aged people is frailty [19] [20], and thus poor physical functioning and overall debilitated condition become important assessments. Several diseases such as hypertension, sarcopenia, cognitive decline and osteoporosis contribute to frailty [21]. As shown by the results, and as frailty scores expectedly rose with age, there were clear sex differences as male rats had a higher average frailty index both at 17- and 27- months. While both aged-male and aged-female rats performed poorly on the treadmill, the females performed better with the grip strength assessment. These sex differences in physical functioning were not reflected in the aged rats when assessed by the open field test. [22]. Both 17- and 27-month-old male and female rats performed poorly on open field testing, which is an indication of general anxiety and an indirect measurement for voluntary movement and threat avoidance. Older rats of both sexes tended to stay away from the center quadrant, while spending the most time in the corners. This age-dependent decline in performance on the open field test suggests that aging increases anxiety behaviors. These results may model findings in human studies where anxiety linked to death and loneliness were higher in older adults [23]. Similarly, anxiety to non-communicable diseases [24] and fear of falling [25] are common in older adults.

Muscle mass can be predictive of physical performance. Loss in muscle mass is often seen as a sign of frailty (i.e., sarcopenia). As seen with the treadmill tests, the older rats performed poorly. Not surprisingly, we observed also an age-dependent decline in muscle weight. Sarcopenia may result from a combination of both loss of muscle mass and increase in fat accumulation in skeletal muscle [26]. To counter or prevent sarcopenia, exercise can potentially help maintain or build muscle, for maintenance during old age[27, 28]. Another study in sarcopenic older adults indicates that while muscle strength improved following exercise programs, the muscle mass did not change in the level of sarcopenia [29]. This may suggest exercise can prevent sarcopenia, but not reverse it.

Body fat content has been used as a surrogate to determine risk to type 2 diabetes mellitus (T2DM)[30–33] And controlling body fat regulates progression of T2DM [34]. Our results show that both male and female rats tend to have higher body fat with old age, but this finding is more pronounced in males. Use of the glucose tolerance test (GTT) can be used to identify T2DM [35, 36]. Aged male rats performed poorly on the GTT, while the aged females were essentially no different from their younger counterparts with normal glucose tolerance.

Measurement of cell cycle inhibitors, *Cdkn2a* (*p16^Ink4a^*) and *Cdkn1a* (*p21*) are few of the several markers used to identify cellular senescence. Clear differences were observed in p*16^Ink4a^* expression (i.e., elevated in femurs of both male and female 27-month-old rats), while no change was seen in the *p21* expression. This elevation in the *p16^Ink4a^*expression correlates well with the decline in femur and spine bone volume fractions, and in the failure to load. The spine BV/TV in the 27-month-old female rats surprisingly did not show this trend to decline at 17 months.

In contrast, both *p16^Ink4a^* and *p21* expression were significantly elevated in the muscle from both male and female aged rats. Together with this, *Gadd45a*, a DNA damage marker, was also elevated significantly in 27-month-old male rats, and non-significantly in the aged female rats. Several SASP markers were also elevated in both male and female aged rats. These increases in senescence and SASP markers correlate well with the decline in *Pax7*, a gene responsible for maintenance of muscle stem cell function, performance on the treadmill, and loss in muscle mass in aged rats.

Interestingly, when senescence markers were assessed in inguinal fat, the 27-month-old male rats had elevated *p16^Ink4a^* and *p21* expression, while there was no change in the corresponding tissue of female aged rats. This data correlates well with the changes in GTT results in aged male rats. These data sets, confirm other studies which have shown that cellular senescence contributes to obesity-related metabolic dysfunction and diabetes[37].

A complete blood cell count is one of the most routinely ordered tests for monitoring during routine health maintenance and for screening of various age-onset and other diseases. Anemia, defined by a decrease in red blood cells or hemoglobin levels, is more common in older adults as compared to younger adults [38]. It has been associated with increased mortality, cognitive impairment and higher incidence of falls and fractures [39–41]. As expected, our results showed a decline in RBC count and Hematocrit values in aged male and female rats. The hemoglobin levels declined with age in females, but not in males. We also observed sex differences in incidence of anemia, as aged female rats where more anemic as compared to aged male rats.

Total white blood cell counts, especially lymphocytes, have been observed to decline with age leading to decreased immunity and increased susceptibility to infections and cancer [42]. In our study we observed a significant decline in total white blood cells and lymphocytes in both aged male and aged female rats. In aging humans and mice there is an hematopoietic stem cell bias towards the myeloid lineage at the expense of lymphopoiesis [43–46], thus the neutrophil counts remain stable with normal aging [46, 47]. Our study also observes no significant change in neutrophil count with aging in rats. Circulating monocyte counts remain stable in older adults [48], but the aged monocyte phenotype is thought to contribute to inflammaging [49, 50]. In our study, while we observe a significant decline in monocyte counts in aged female but not in male rats.

Aging is associated with a decline in platelet counts however there is an increase in platelet aggregability which leads to increased incidence of thrombotic events in older adults. We observed a decline in platelet count with age in both male and female rats, with aged female rats showing a lower platelet count compared to aged males.

In summary, our data show that rats can be used as an alternate, reliable and easy to use model to study aging and to assess efficacy of interventions targeted to ameliorate age-related morbidities. While this manuscript is limited in scope comparing the advantages of using rats versus other non-human primates, these data sets provide a reference point for future pre-clinical studies done in rats, especially studies done in aged genetic models of rats. In summary, the study of rat physiology provides a valuable foundation for understanding fundamental biological processes and disease mechanisms. While rats share many physiological similarities with humans, notable differences in metabolism, immune response, and organ function must be carefully considered when extrapolating findings.

## Materials and Methods

### Animal studies

Animal studies were approved by the Institutional Animal Care and Use Committee at Mayo Clinic. Brown-Norway rats were housed in our facility at 23°C to 25°C with a 12-hour light/ dark cycle and were fed with standard laboratory chow (Teklad Global 14% Protein Rodent Maintenance Diet, Envigo, Madison WI, USA) with free access to water.

### Frailty

Frailty index [33] scoring in rats was used to assess andt quantify the physical decline and health status of rats in the context of longitudinal aging. Each component was assigned a numerical score based on the rat’s performance or appearance as described previously [13]. Using a 30-point scoring which included changes in the integument, physical or musculoskeletal changes, auditory changes, ocular or nasal changes, and urogenital changes, the rats when scored for a noticeable change got a score of 1, while less noticeable changes got a score of 0.5 and no indication or well-conditioned got a score of 0.

### Grip Strength Test

Rats were grasped by the base of the tail and the scruff of the neck and suspended above a grip ring. The rats were gently lowered toward the grip ring and allowed to grasp the ring with its forepaws. The rats were lowered to a horizontal position and the tail was tugged until they released the ring, the body being supported by grasping the scruff of the neck. Forelimb grip strength (NF/kg) was determined using a Grip Strength Meter from Columbus Instruments, OH. The mean force was determined with a computerized electronic pull strain gauge that is fitted directly to the grasping ring and the force was normalized to body weight. Average measurements from three successful trials were used for calculations.

### Open Field Test

The open field test was used to evaluate the sensorimotor parameters associated with general activity levels, gross locomotor activity, and exploration habits. The degree of activity in the open field was measured by the number of times the animal moves into the various marked off areas. Typically, rodents exhibit anxiety-like behaviors by preferring the edges of the arena (periphery) over the center. Spending more time in the center can be interpreted as reduced anxiety. Rats were assessed in sound-insulated rectangular activity chambers from TSE Systems, On the test day, following acclimatization for 10 minutes rats were introduced into the chambers.

### Treadmill test

Rats were first acclimated to the treadmill for 3 consecutive days for a duration of 6 minutes on each day at a speed of 3m/min for 2 min, 4m/min for 2 min and 5m/min for 2 min at a 5° incline (Columbus Instruments). On the test day, the rats ran on the treadmill at an initial speed of 5 m/min for 2 min, and then the speed was increased by 2 m/min every 2 min until the rats were exhausted. Exhaustion was defined as the inability to return onto the treadmill despite a mild electrical shock stimulus and mechanical prodding. Total distance travelled and treadmill speed were recorded.

### Glucose Tolerance test

Rats were fasted for 6 hours before the GTT. The rats were then given 1.0 mg/kg of glucose by oral gavage. Blood was collected through a nail trim method. For the GTT, 2μl of blood was collected, and glucose was measured using a handheld glucometer at time 0, 15-, 30-, 60- and 120- min.

### Quantitative Real Time PCR

Fat, muscle, and bone samples were collected for mRNA isolation and qRT-PCR. The samples were homogenized, and total RNA was isolated using RNeasy Mini Columns (QIAGEN, Valencia, CA). cDNA was generated from mRNA using the High-Capacity cDNA Reverse Transcription Kit (Applied Biosystems by Life Technologies, Foster City, CA) according to the manufacturer’s instructions, and RT-qPCR was performed as described in our previous studies[51]. All primer sequences have been authenticated in previous studies. Primers were designed so that they overlapped two exons. A detailed list of primer sequences is provided in Tables S1 and S2.

### Bone architecture assessment

Bone architecture assessments were done using viva CT 40. Femur and Spine were scanned at 10.5 µm scanned using microCT (vivaCT 40, Scanco Medical AG,Brüttisellen, Switzerland). The distal femur was scanned corresponding to a 1-5 mm area above the growth plate. All images were first smoothed by a Gaussian filter (sigma=1.2, support=2.0), then thresholded corresponding to 30% of the maximum available gray scale values. Volumetric bone mineral density (vBMD), bone volume fraction (BV/TV), trabecular thickness (Tb.Th), trabecular separation (Tb.Sp), trabecular number (Tb.N), and structure-model index (SMI) were calculated using 3D standard microstructural analysis.

### Complete blood count and blood chemistry assessment

Whole blood was collected in Greiner Bio-One MiniCollect Tubes with K_2_ EDTA (Ref# 45048) and mixed thoroughly by inverting the tubes multiple times. The samples were analyzed by the Department of Molecular Medicine core at Mayo Clinic, on an Abaxis VetScan HM5 Analyzer for the detection of complete blood count. Blood chemistry assay was performed on the Piccolo^®^ Xpress™ Chemistry Analyzer. For this the blood was collected in the Greiner Bio-One Minicollect Tubes with Lithium Heparin (Ref#450477) and analyzed for various parameters.

### Statistical analysis

The analysis and sample size calculations were facilitated by the biostatistics core at the Mayo Clinic. All statistics were performed using GraphPad Prism 8.1.1 software. Data are expressed as median with interquartile range and analyzed by an unpaired two-tailed t-test when comparing two groups and one variable, a one-way ANOVA, when comparing three groups or a two-way ANOVA, when comparing two or more groups and variables.

## Acknowledgements

This work was made possible by the Robert and Arlene Kogod Professorship (to RJP); P01 AG062413 (SK, JNF, RJP, JLK, TT), R01 AG063707 (DGM), R01 R01AG068048 (JFP), 1UG3CA26810-01 (JFP), R01AG063543 (LJN), U19AG056278 (PDR/ LJN), R01 AG082681 (A.C), R01 DK128552 (JNF), and K01 AR070241 (JNF). We would like to thank Christine Hachfeld, Jennifer Hager, and Claire Wilhelm for their technical help with animal husbandry and behavioral testing.

## Author Contributions

AC and RJP designed the study, AC, XZ, JNF and AD performed the experiments, while AC, DGM, JNF, AD, RS JP, RJP and SK analyzed and interpreted the data. AC and RJP wrote the initial manuscript, and all authors contributed ideas and revised and approved the final version of the manuscript.

## Data Availability

All data will be made available on request.

---

**Supplementary Figure 1.**
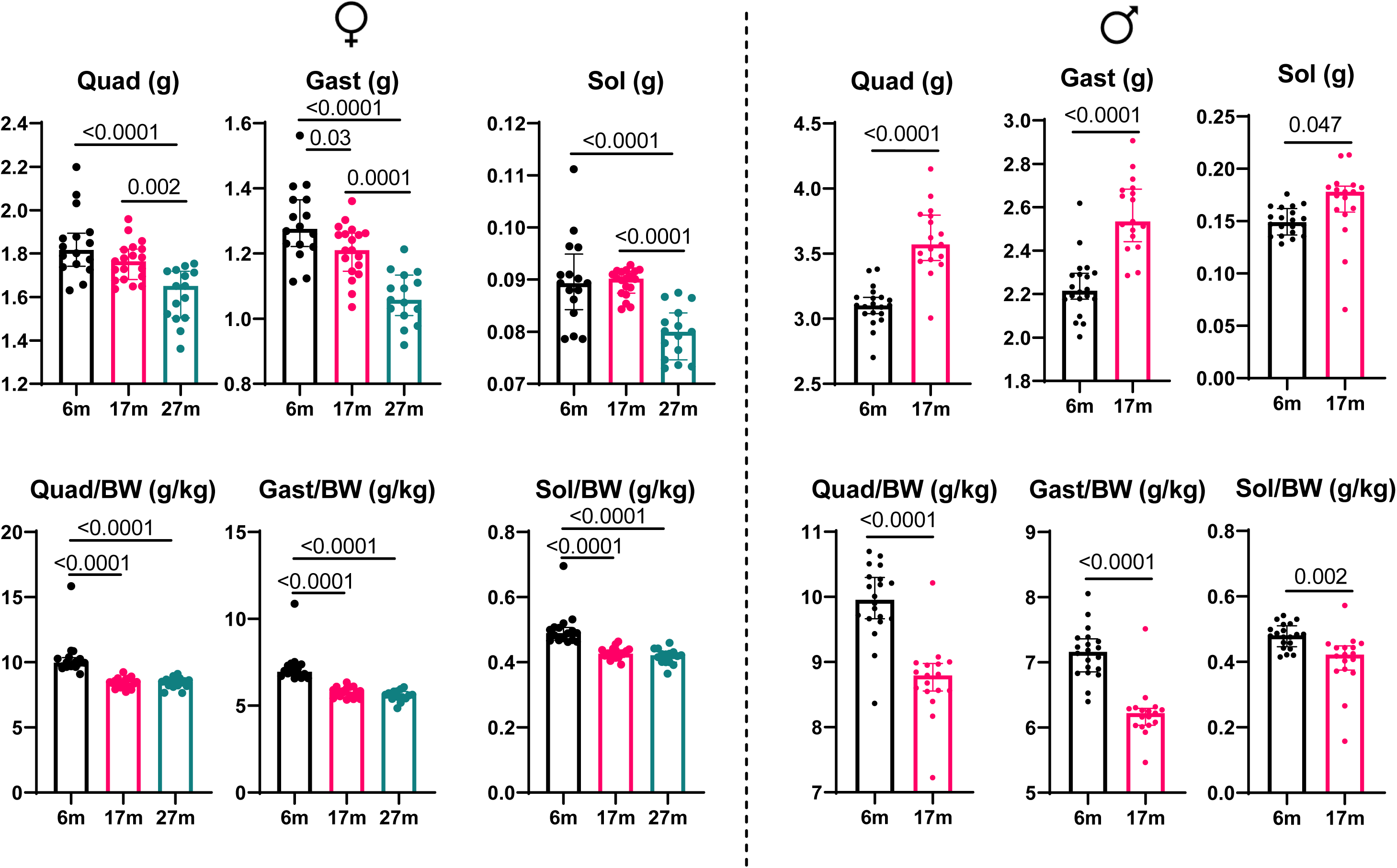
Age-related changes in muscle weight. Muscle weights for quadriceps, gastrocnemius and soleus were measured in male and female Brown Norway rats. Statistical analyses were done using GraphPad Prism and p values were calculated using One-way ANOVA for females and two-tailed t-test for males.

**Supplementary Figure 2.**
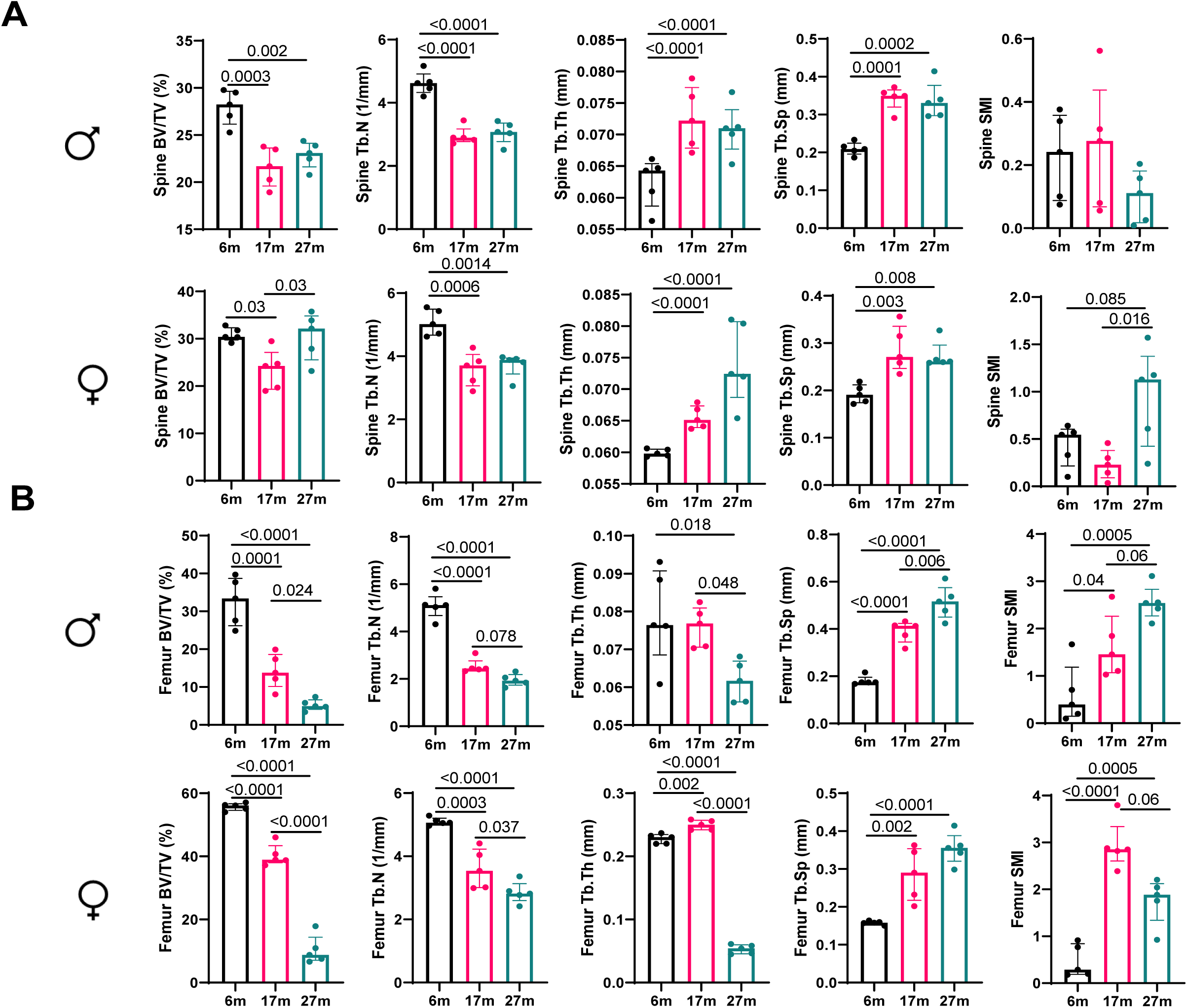
Age-related changes in bone parameters in rats. Micro-CT parameters for spine (A) and femur (B) are shown in 6-, 17- and 27-month-old male and female Brown Norway rats. Statistical analyses were done using GraphPad Prism and p values were calculated using One-way ANOVA.

**Supplementary Figure 3.**
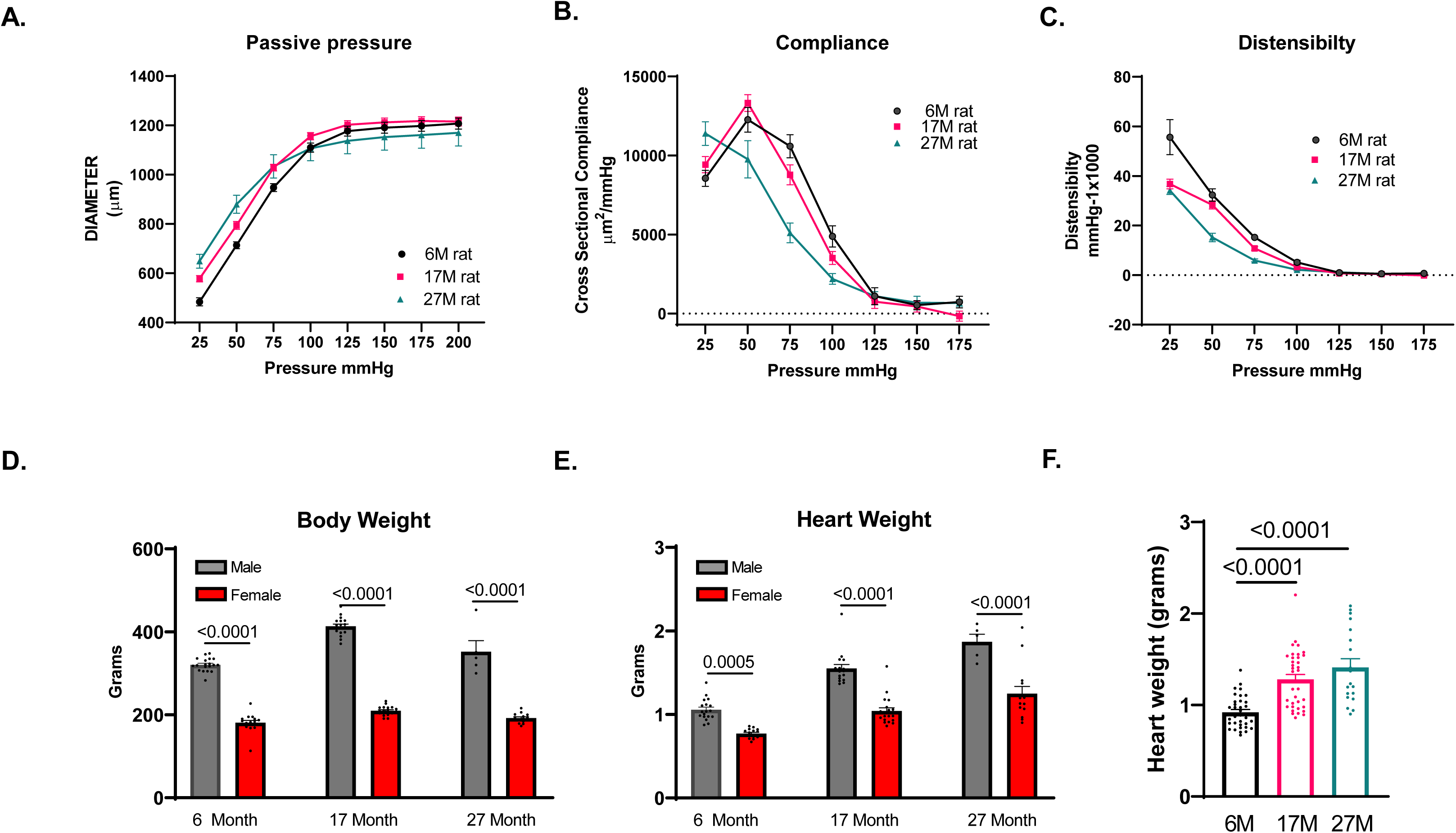
Age-related changes in heart. Heart functions were assessed using passive pressure (A), compliance (B) and distensibility (C) with no significant changes seen between 6-, 17- and 27-month-old Brown Norway rats. Body weight (D) and heart weight (E) for 6-, 17- and 27-month-old male and female Brown Norway rats are presented. (F) Heart weight for both male and female 6-, 17- and 27-month-old Brown Norway rats are presented.

**Table S1:**
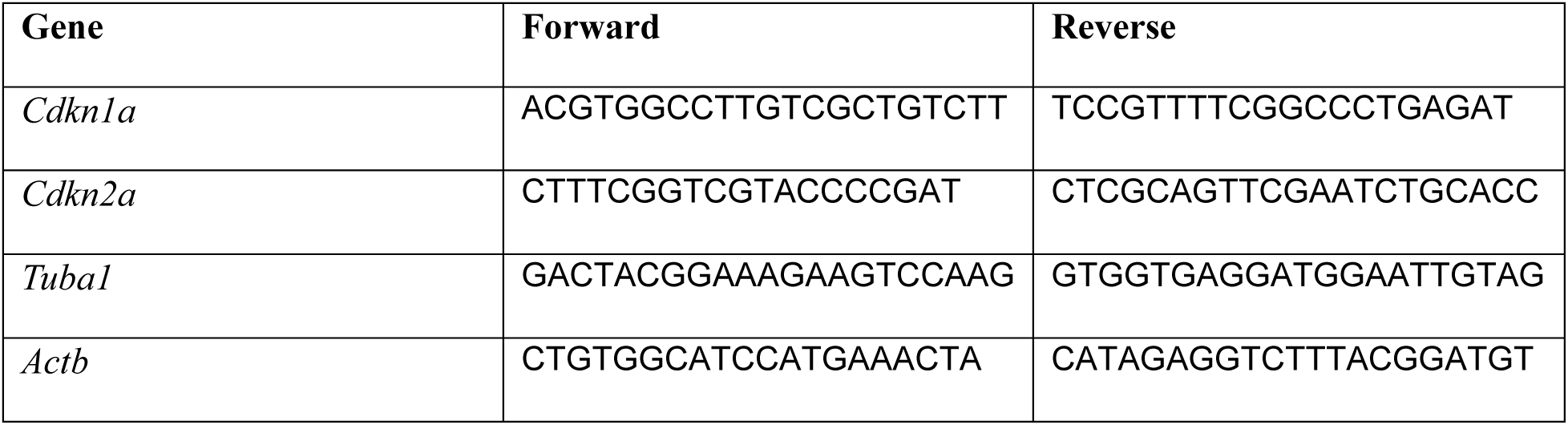
Rat qPCR primers in figures 4.

**Table S2:** Rat qPCR primers in figure 4 and 5.

| **Gene** | **Source** | **Identifier** |
| --- | --- | --- |
| *Cd68* | Integrated DNA Technologies | Rn.PT.58.37733352 |
| *Cdkn1a* | Applied Biosystems | Rn00589996_m1 |
| *Cdkn2a* | Applied Biosystems | Rn00580664_m1 |
| *Gadd45a* | Integrated DNA Technologies | Rn.PT.58.7466023 |
| *Tnf* | Integrated DNA Technologies | Rn.PT.58.11142874 |
| *Cxcl10* | *Integrated DNA Technologies* | *Rn.PT.58.35118131* |

